# Glycerolipid defects in skeletal muscle contribute to rhabdomyolysis in Tango2 deficiency

**DOI:** 10.1101/2022.11.12.516274

**Authors:** Jennifer G Casey, Euri S Kim, Brian S Tao, Arian Mansur, E. Diane Wallace, Vandana A Gupta

## Abstract

Rhabdomyolysis is a clinical emergency characterized by severe muscle damage resulting in the release of intracellular muscle components leading to myoglobinuria and in severe cases, acute kidney failure. Rhabdomyolysis is caused by genetic factors that are linked to increased disease susceptibility in response to extrinsic triggers. Recessive mutations in *TANGO2* result in episodic rhabdomyolysis, metabolic crises, encephalopathy and cardiac arrhythmia, the underlying mechanism contributing to disease onset in response to specific triggers remains unclear. To address these challenges, we created a zebrafish model of Tango2 deficiency. Here we show that loss of Tango2 in zebrafish results in growth defects, early lethality and increased susceptibility of muscle defects similar to *TANGO2* patients. Detailed analyses of skeletal muscle revealed defects in the sarcoplasmic reticulum and mitochondria at the onset of disease development. The sarcoplasmic reticulum (SR) constitutes the primary lipid biosynthesis site and regulates calcium handling in skeletal muscle to control excitation-contraction coupling. Tango2 deficient SR exhibits increased sensitivity to calcium release that was partly restored by inhibition of Ryr1-mediated Ca^2+^ release in skeletal muscle. Using lipidomics, we identified alterations in the glycerolipid state of *tango2* mutants which is critical for membrane stability and energy balance. Therefore, these studies provide insight into key disease processes in Tango2 deficiency and increased our understanding of how specific defects can predispose to environmental triggers in TANGO2-related disorders.

## Introduction

Rhabdomyolysis (RM) is a complex medical disorder involving catastrophic failure of skeletal muscle homeostasis and integrity, resulting in muscle breakdown and release of muscle cytosolic content into circulation (Cabrera-Serrano & Ravenscroft, 2022). Rhabdomyolysis can give rise to serious health complications such as myoglobinuria, cardiac arrhythmia and acute kidney injury. Clinical symptoms can include severe muscle weakness, myalgia, and muscle swelling with a serum creatine kinase (CK) rising above 1000U/l. Rare disease-causing mutations are associated with a small but significant subset (∼15%) of RM patients. Environmental factors such as viral infections (SARS-CoV-2, HIV), physical exertion, certain medications and drugs are major contributing factors in combination with a genetic predisposition (East, Alivizatos, Grundy, Jones, & Farmer, 1988; Knoblauch, Dagnino-Acosta, & Hamilton, 2013; Rawson, Clarkson, & Tarnopolsky, 2017; Szugye, 2020; van den Bersselaar et al., 2021; Wu, Wong, Cheng, & Yu, 2022). Even in genetic forms of RM, environmental factors increase the susceptibility to recurrent episodes of muscle breakdown (Kruijt et al., 2021). Environmental factors contributing to RM have been mostly identified through life-threatening reactions to different triggers in the clinical population. Lack of clear understanding of intrinsic disease mechanism also prevents investigating the role of extrinsic factors on increasing the susceptibility of muscle damage in predisposing genotypes.

Recessive mutations in *TANGO2* underlie a rare pediatric disorder resulting in encephalopathy, rhabdomyolysis and cardiac abnormalities (Kremer et al., 2016; Lalani et al., 2016; Miyake et al., 2022). *TANGO2* encodes for “Transport and Golgi Organization 2” protein, first identified in a genetic screen for proteins required for constitutive protein secretion in *Drosophila* cells (Bard et al., 2006). Depletion of Tango2 results in the fusion of endoplasmic reticulum and Golgi compartments in *Drosophila* cells. TANGO2 deficient patients’ fibroblasts exhibit a profound decrease in the endoplasmic reticulum area (Lalani et al., 2016). Functional studies in patients’ fibroblasts have shown that TANGO2 is required for ER-Golgi trafficking in cells (Milev et al., 2021). Proteomic analysis of fibroblasts from *TANGO2* patients revealed significant changes in the component of mitochondrial fatty acid oxidation, plasma membrane, endoplasmic reticulum, Golgi and secretory pathway indicating pleiotropic roles in disease biology (Mingirulli et al., 2020). Some patients’ fibroblasts also showed a defect in palmitate-dependent oxygen consumption suggestive of impaired mitochondrial fatty acid oxidation (Heiman et al., 2022; Kremer et al., 2016). In contrast, myoblasts from TANGO2 patients exhibit no defects in mitochondrial structure and function but exhibit abnormalities in mitochondrial function under nutrient stress suggesting the involvement of extrinsic factors in inducing cellular defects (Bérat et al., 2021). These studies suggest TANGO2 deficiency results in intrinsic metabolic defects that are exacerbated under stress conditions, but a clear understanding of the disease processes resulting in the pathological state is lacking.

*TANGO2* mutations result in clinical heterogeneity in affected patients. Muscle weakness and neurodevelopmental presentation precede life-threatening complications of rhabdomyolysis and cardiac arrhythmias or cardiomyopathy in many cases. However, a clear genotype-phenotype correlation is lacking in these patients (Powell, Ames, Knierbein, Hannibal, & Mackenzie, 2021). The presence of variable phenotypes in different cell types suggests TANGO2 may play diverse roles in various cell types *in vivo. Tango2* knockout mice exhibit normal development, lifespan and physiology (International Mouse Phenotyping Consortium). Therefore, robust model systems are needed to understand the effect of Tango2 deficiency on variable clinical presentation and identify the pathological processes contributing to disease onset and progression. To address these challenges, we have developed vertebrate zebrafish models of Tango2 deficiency. Tango2 deficiency results in normal embryonic development in zebrafish but causes increased lethality during larval and juvenile stages. *tango2* mutant larval zebrafish exhibit sarcoplasmic reticulum and mitochondria defects and develop acute muscle dysfunction on triggering sarcoplasmic reticulum stress. Global lipidomic identified a reduced abundance of lipids synthesized by ER/SR localized enzymes in Tango2 deficiency which are critical for cellular membranes and energy states. The studies presented in this work provide mechanistic insights into intrinsic disease processes in *TANGO2*-related disorders.

## Results

### Tango2 deficiency in zebrafish results in normal embryonic development but increased lethality during larval and juvenile stages

Zebrafish grow *ex vivo* and therefore disease onset and progression can be visualized in individual animals. To understand the role of Tango2 *in vivo* in disease onset and progression, we created the loss of function *tango2* alleles in zebrafish using the CRISPR-Cas9 system. *tango2* gene in zebrafish encodes for two transcripts and therefore sgRNAs were designed to knock out both transcripts (Figure 1A). *tango2* alleles created include *tango2*^bwg210^ with insertion of seven bases (c.226_227ins7; p.Tyr76Leufs*25) and *tango2*^bwg211^ with 26 base insertions in exon2 (c.226_227ins26; p.Tyr76Leufs*207) (Figure 1B). These alleles result in out of frame mutations and are predicted to result in truncating proteins. Phenotypic analysis of *tango2*^bwg210^ and *tango2*^bwg211^ lines revealed no significant differences, therefore, *tango2*^bwg211^ line (referred as *tango2* mutants) was used for further investigation.

**Figure 1.**
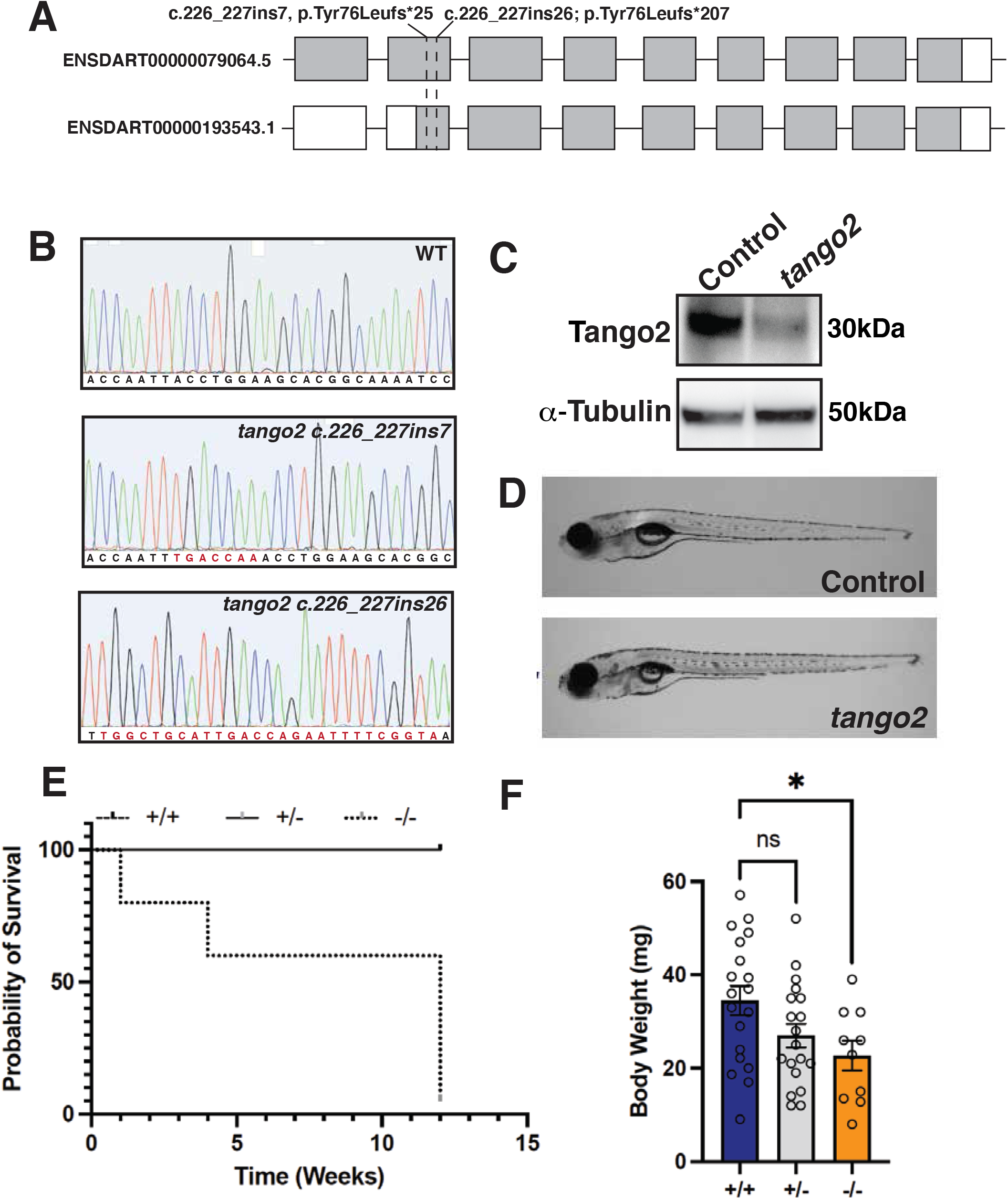
Tango2 deficiency results in growth defects and lethality. (A) Schematics depicting the position of *tango2* alleles (*tango2*^*bwh210*^ and *tango2*^*bwh211*^) with reference to both transcripts in zebrafish. *tango2* mutations resulted in insertions and frameshift mutations in different alleles. (B) Sanger sequencing for control and *tango2* alleles depicting mutations in *tango2*^*bwh210*^ (c.226_227ins7) and *tango2*^*bwh21*^ (c.226_227ins26). (C) Western blot analysis of protein from control and *tango2* mutant fish (1 month age). (D) Lateral view of control and *tango2* mutant larval zebrafish showing no phenotypic differences. (E) Kaplan-Meier survival curve of control (+/+ and +/-) and *tango2* mutants (-/-); n=360. (F) Body weight analysis of control (+/+ and +/-) and *tango2* mutants (-/-) at 1 month; n=60 and data are presented as mean±S.E.M (unpaired t-test, parametric); *p<0.05.

To validate the effect of c.226_227ins26 mutation on protein level, Western blot analysis was performed in control and *tango2* zebrafish (4 weeks) that showed a complete absence of tango2 protein in the mutant fish (Figure 1C). Therefore, c.226_227ins26 mutation results in the loss of function of Tango2 protein in zebrafish. Phenotypic analysis of control and *tango2* mutant larval zebrafish showed no significant morphological differences during early development (Figure 1D, 8 days post fertilization; dpf). To identify any effect of tango2 deficiency on the lifespan of mutant fish, +/+, +/- and -/-genotypes were analyzed till 3 months of age. Despite no obvious morphological differences in control (+/+) and mutant (-/-) larval fish at early stages, a reduced survival was observed for mutant fish from 7 days post fertilization and most of the mutant fish died by 3 months (4% survival) (Figure 1E). Quantification of body weight also showed a significant reduction in body weights of *tango2* mutants in comparison to wild-type siblings at four weeks of age indicative of growth defects (Figure 1F). Together, these studies suggest that Tango2 deficiency in zebrafish results in growth defects and early lethality in larval and juvenile stages.

### Tango2 is localized at endo-membranes in the I-band region

*TANGO2* mutations result in a decreased network of the endoplasmic reticulum in patients’ fibroblasts and metabolic abnormalities indicative of involvement of these intracellular organelles with disease pathology. To address the localization of Tango2 in skeletal muscle, immunofluorescence was performed on myofibers isolated from zebrafish (8 weeks). Co-localization with different skeletal muscle proteins showed partial co-localization of Tango2 with Ryr1 (sarcoplasmic reticulum protein), Tomm20 (mitochondrial protein) and Golga2 (Golgi apparatus) in the I-band-Zdisc region (alpha-actinin). Analysis of *tango2* mutant myofibers showed disorganized regions in a few myofibers (∼10%) by Ryr1 staining indicative of structural defects in SR in skeletal muscle. No changes were observed in Golga2 or alpha-actinin. Therefore, these studies suggest that Tango2 is localized and regulates the structure of sarcoplasmic reticulum and mitochondria.

### Tango2 mutant fish exhibit ultrastructural defects in the sarcoplasmic reticulum and mitochondria at disease onset

To identify the temporal onset of structural abnormalities observed in myofibers at larval fish (45 days, Figure 2), skeletal muscle ultrastructure was examined during early larval development (8 dpf) when control and mutant fish are phenotypically and functionally similar. Ultrastructural evaluation by electron microscopy showed no significant defects in sarcomere size (length or height) in *tango* mutants in comparison to control siblings (Figure 3). While most of the sarcoplasmic reticulum and mitochondria were normal in *tango2* mutants, few myofibers exhibited defects in sarcoplasmic reticulum and mitochondria structures (Figure 3). Interestingly, both of these abnormalities were present together in the affected myofibers and no myofibers with either sarcoplasmic reticulum or mitochondrial defects were observed. Defective sarcoplasmic reticulum showed either collapsed or smaller terminal cisternae of sarcoplasmic reticulum in *tango2* mutants in comparison to controls (Figure 3, arrows). Moreover, an abnormal accumulation of vesicles was also observed in the proximity of the sarcoplasmic reticulum. In comparison to longer mitochondria in control muscles, *tango2* mutant muscle exhibited smaller mitochondria associated with whorled membrane structures that are also seen in mitochondrial myopathies (Figure 3, M) (Vincent et al., 2016). These data show that the absence of Tango2 results in ultrastructure defects in the sarcoplasmic reticulum and mitochondria during the pre-symptomatic disease state (8 dpf).

**Figure 2.**
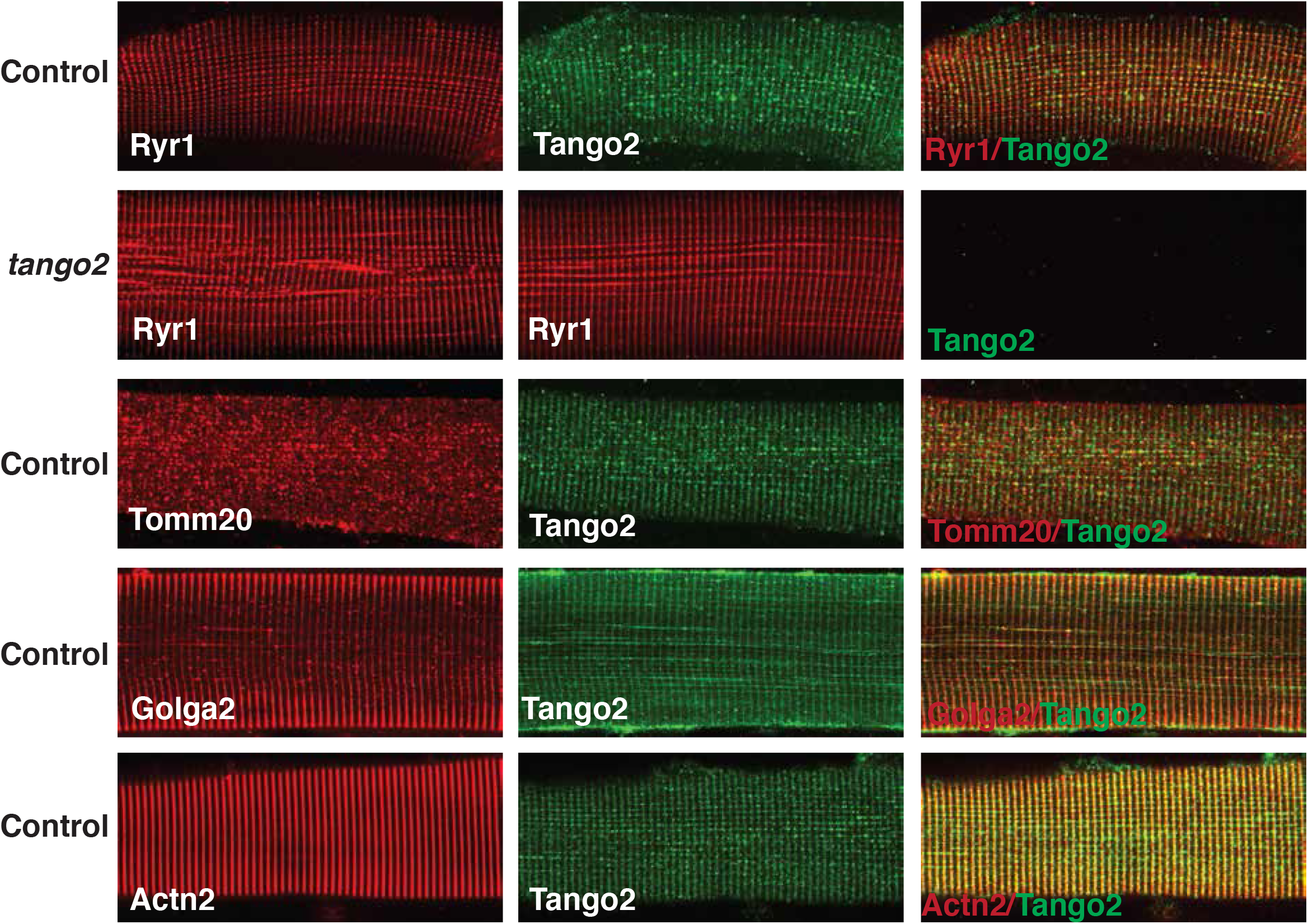
Tango2 is localized to SR, mitochondria and Golgi in the I-band region. Myofibers were isolated from control or *tango2* mutants (1.5 months) zebrafish and immunofluorescence was performed. Myofibers were isolated from 3 different zebrafish for each genotype and 10-15 myofibers were analyzed in each group.

**Figure 3.**
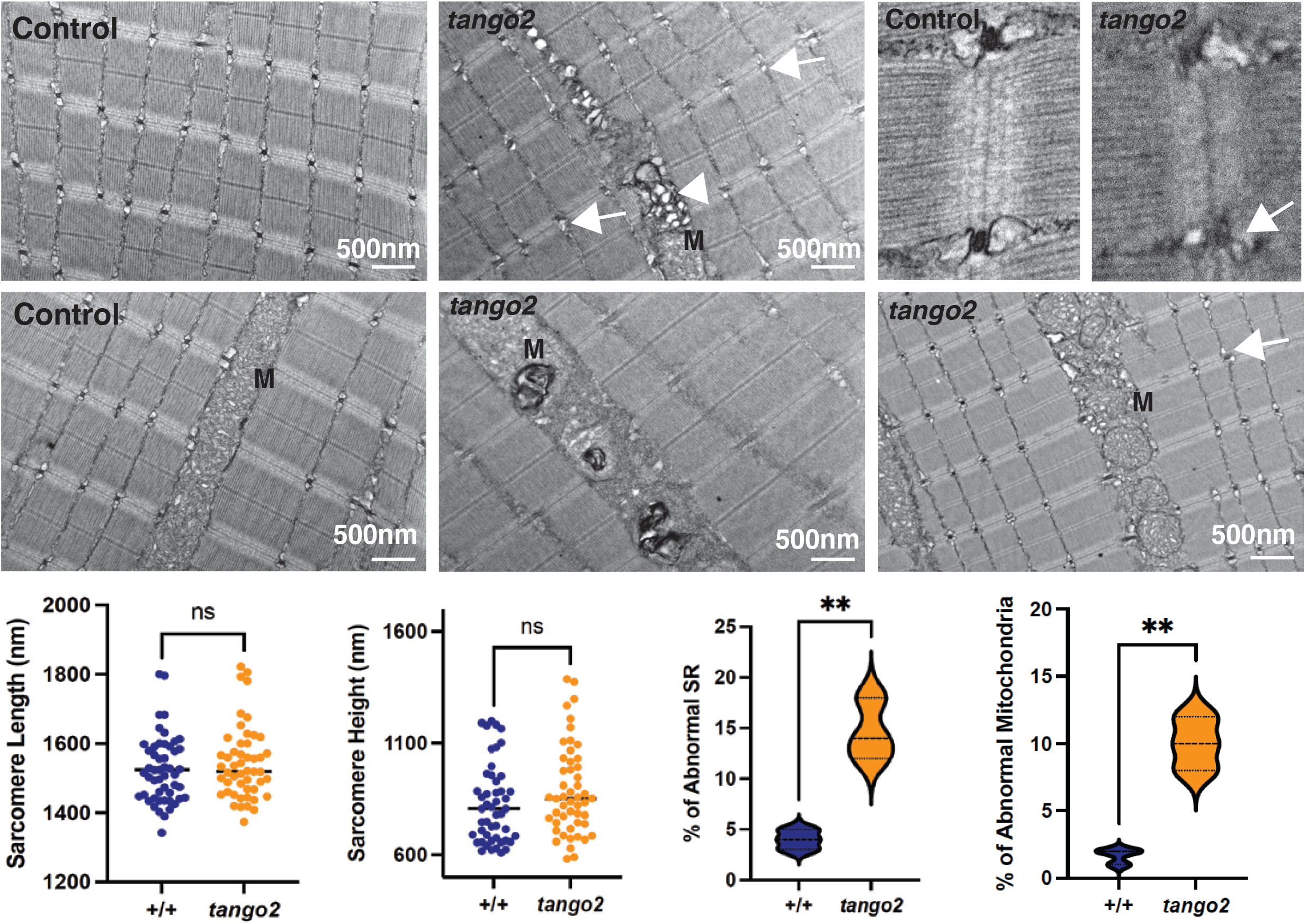
Tango2 deficiency results in the abnormal sarcoplasmic reticulum and mitochondrial in skeletal muscle. Transmission electron microcopy was performed on the longitudinal sections of control and *tango2* mutant larval fish (8 dpf). *tango2* mutants display normal sarcomere size compared to controls. A small number of sarcoplasmic reticulum exhibited small or collapsed sarcoplasmic reticulum (arrows) and accumulation of abnormal vesicles (arrow). Some of the mitochondria in *tango2* mutants also exhibited abnormal mitochondria (M) with whorled membrane structures or small size in comparison to controls. Data represent mean±S.E.M from three different samples. Non-significant: ns, \*\**p*<0.01, Student’s t test.

### SR stress results in acute motor dysfunction in *tango2* mutants

*tango2* mutant larvae exhibit normal motor function during early larval stages in spite of ultra-structural changes in a small group of myofibers, (5-7 dpf, Figure 4). This is similar to many *TANGO2* patients that exhibit normal motor function except for developing acute muscle damage during rhabdomyolysis episodes. *tango2* mutant fish exhibit defects in the terminal cisternae of the sarcoplasmic reticulum that is occupied by Ryr1 channels (Figure 2). SR is the regulator of excitation-contraction coupling in skeletal muscle through release of Ca^2+^ by the Ryr1 channel and reuptake of Ca^2+^ by the Serca channel (Lawal et al., 2020). Caffeine is an activator of Ryr1 that binds to Ryr1 and increases Ca^2+^ sensitivity. To test if tango2 deficiency results in increased susceptibility to muscle damage caused by sarcoplasmic reticulum stress, we treated control and *tango2* mutant larval fish (6 dpf) with caffeine and quantified the swimming behavior by using an automated movement tracking system. Acute caffeine exposure resulted in reduced swimming of controls (+/+, +/-) as well as *tango2* mutant (-/-) fish, but the effect was more severe in *tango2* mutants in comparison to control siblings. While control fish recovered completely after 24 hours of caffeine exposure, *tango2* mutants failed to revert to the normal level of motor function. This suggests that tango2 deficiency results in increased susceptibility to muscle damage in response to extrinsic triggers affecting SR function.

**Figure 4.**
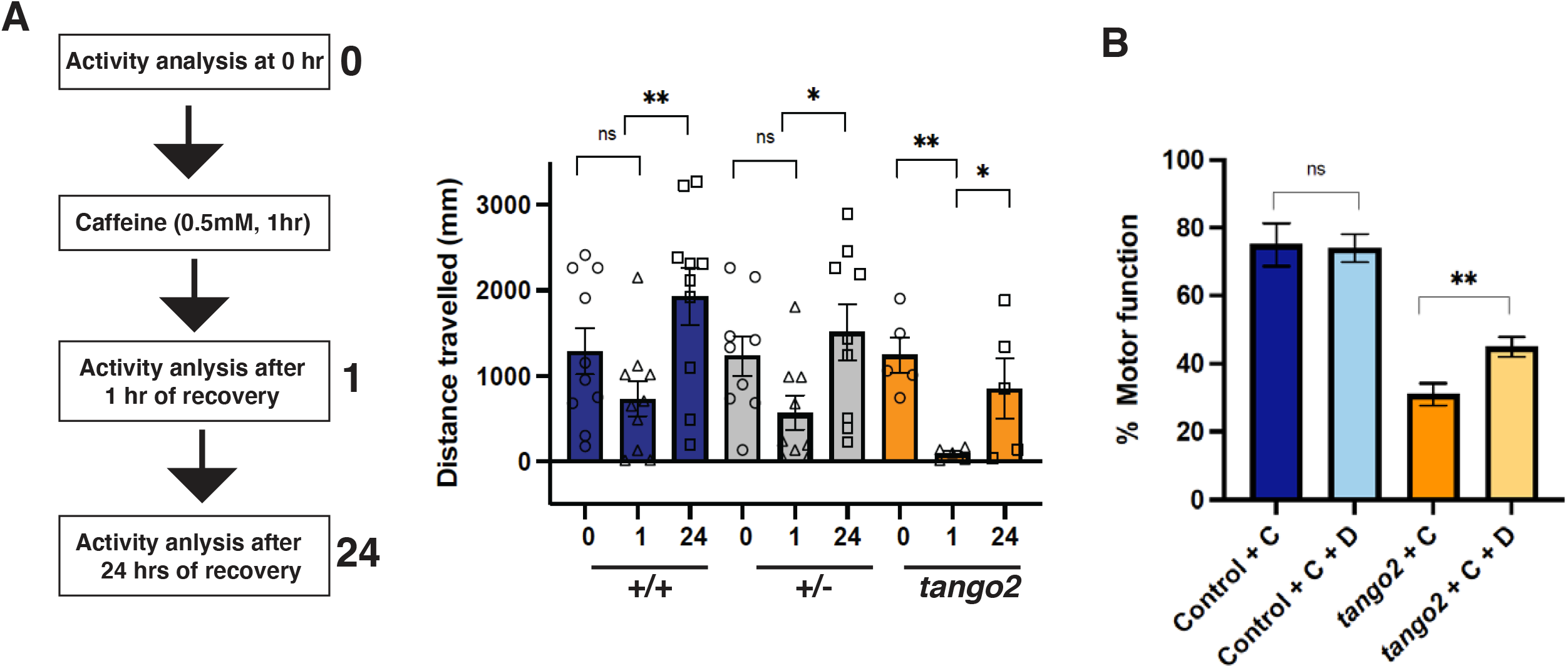
*Tango2* mutants exhibit increased sensitivity to caffeine-induced SR stress that is partially reversed by dantrolene. (A) Schematics for caffeine treatment and effect of caffeine treatment on total swimming distance by control (+/+ and +/-) and *tango2* mutants (-/-); n=96 (B) Effect of dantrolene on motor function in caffeine treated control (+/+ and +/-) and *tango2* mutants (-/-). Data was normalized to swimming distance before caffeine treatment within each group. N=48 in each group with 10-22 of different genotypes. Data represent mean±S.E.M. Non-significant: ns, \**p*<0.05, \*\**p*<0.01, Student’s t test.

### Dantrolene partially restores caffeine-induced motor deficits in *tango2* mutants

Dantrolene is a Ryr1 antagonist and protects against hypersensitivity of calcium release from the sarcoplasmic reticulum and is used clinically to control malignant hyperthermia and rhabdomyolysis. Therefore, we tested the efficacy of dantrolene in improving acute muscle dysfunction in *tango2* mutants induced by caffeine exposure. Control and *tango2* mutant larval zebrafish (6 dpf) were treated with caffeine (C) or caffeine and dantrolene (D) and normalized motor function (to pre-caffeine treatment) was analyzed. Caffeine resulted in a highly reduced motor function in *tango2* larval zebrafish in comparison to controls (Figure 4B). Treatment with dantrolene resulted in a small improvement in muscle function in *tango2* mutants. Although an improvement in motor function was observed in *tango2* mutants, dantrolene treatment did not result in a complete rescue of muscle function to normal levels.

### Altered lipids profiles in Tango2 deficiency

The majority of cellular lipids are synthesized in ER/SR which is the central hub to regulate cellular lipids composition in repones to intrinsic, homeostatic and environmental factors. Fibroblasts from *TANGO2* patients exhibit abnormal accumulation of fatty acids, however, a clear correlation of lipids in disease pathology is not established due to clinical heterogeneity in patients’ samples. Therefore, to comprehensively characterize the effect of Tango2 deficiency on the content and composition of structural lipids, lipidomics was performed in control and *tango2* mutants (4 weeks) in absence of any extrinsic trigger (Figure 5A). *tango2* mutants revealed a significant decrease in the abundance of phosphatidylcholine (PC) and triglycerides with a small decrease in phosphatidylcholine (PE) (Figure 5B). PC is metabolized to lysophosphatidylcholine (LPC) and free fatty acids. Reduced levels of LPC were also observed in *tango2* mutants suggesting a low abundance of PC results in decreased levels of downstream lipids such as LPC in tango2 deficiency. No significant changes were seen in other lipid classes in Tango2 deficiency. ER/SR harbors enzymes for the glycerol-3-phosphate pathway that synthesizes phospholipids which are major building blocks of lipids for cellular membrane. Therefore, the overall abundance of major membrane and cellular lipids synthesized through ER/SR is significantly decreased in *tango2* mutant zebrafish at the basal level in the absence of any external trigger.

**Figure 5.**
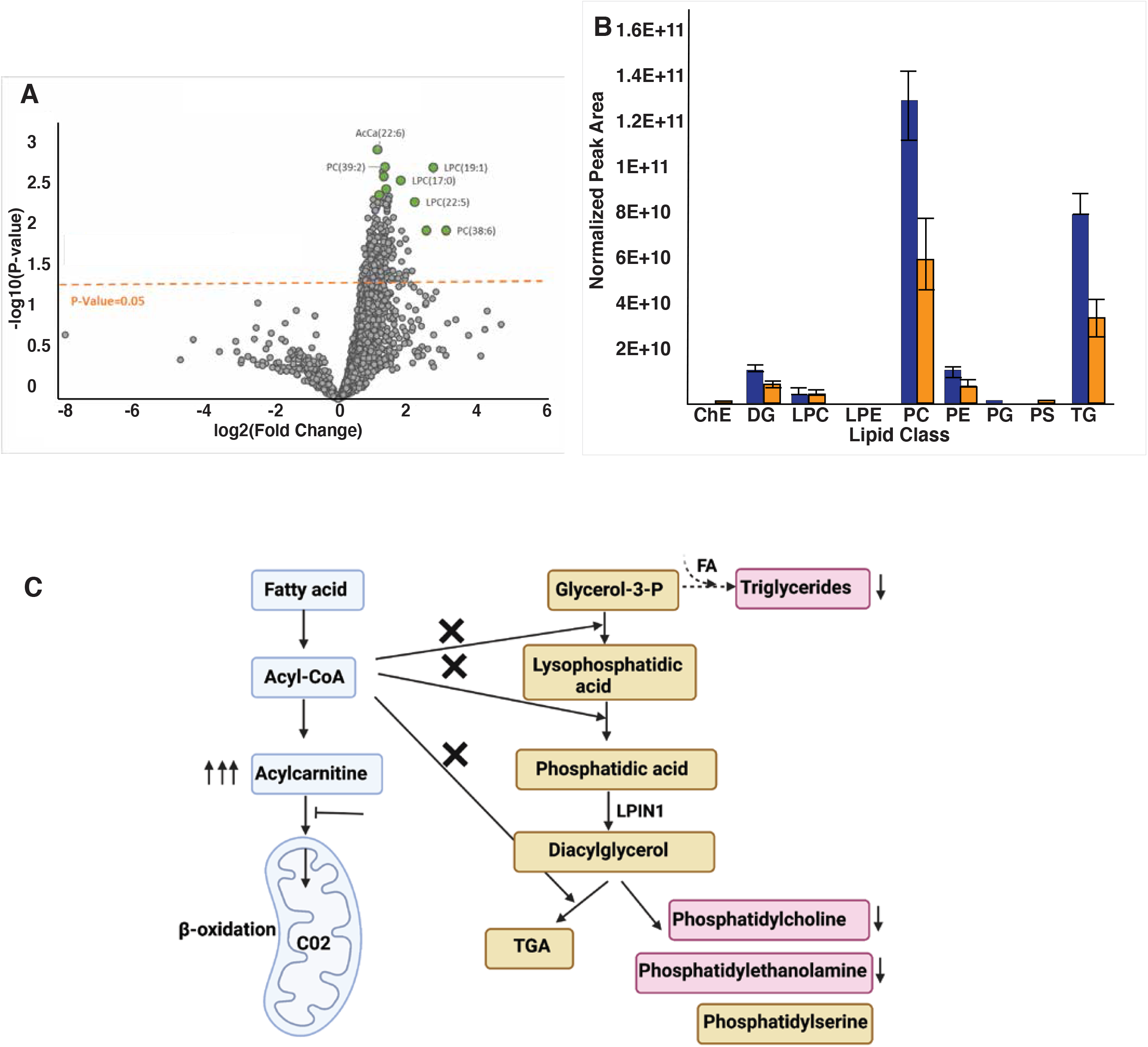
Changes in lipid composition in Tango2 deficiency. (A) Volcano plot of the lipids fold change in control vs *tango2* mutants (4 weeks). Fold differences are calculated as the ratio of lipids in WT/t*ango2* mutants. Dots in green represent the lipids with the lowest levels in *tango2* mutants in comparison to controls (B) Normalized profiles of different lipids classes (Blue: control, Orange: *tango2* mutants) (C) Working model of the effect of Tango2 deficiency on lipid metabolism: *tango2* mutants exhibit decreased abundance of phospholipids and triglycerides (Pink) indicating a defect in the glycerol-3-P pathway. Protein product of *LPIN1*, a rhabdomyolysis causing gene, catalyzes diacylglycerol synthesis in the glycerol-3-P pathway. Triglycerides are energy stores which are synthesized utilizing glycerol-3-phosphate and fatty acids (FA) and reduced in Tango2 deficiency. Some patients’ fibroblasts exhibit an accumulation of acylcarnitine which may limit the availability of acyl-CoA for the glycerol-3-P pathway thereby resulting in phospholipid deficiency. TGA: Triacylglycerol; ChE: Cholesterol ester; DG: Diglyceride ; LPC: Lysophosphatidylcholine; LPE: Lysophosphatidylethanolamine; PC: Phosphatidylcholine; PE: Phosphatidylethanolamine; PG: Phosphatidylglycerol; PS: Phosphatidylserine; TG: Triglyceride.

## Discussion

Rhabdomyolysis is a complex condition with several clinical complications and entails the rapid dissolution of damaged skeletal muscle, often leading to a life-threatening condition. Rhabdomyolysis state is a combination of environmental factors such as infections, fasting, drugs, medications, heat or other unknown triggers and predisposing genotypes. However, a lack of a clear understanding of the intrinsic processes that contribute to disease pathology precludes the assessment of different triggers and their interaction with different genetic susceptibilities in rhabdomyolysis. To address these questions, we developed a knockout zebrafish model of Tango2 deficiency that recapitulates functional and pathological changes observed in *TANGO2* patients and provide the mechanism for skeletal muscle defects.

Our *tango2* zebrafish showed normal embryonic development and motor function during larval stages. This is similar to TANGO2 patients that typically do not exhibit early embryonic developmental defects and develop metabolic crisis and rhabdomyolysis during early or late childhood. Therefore, the *tango2* zebrafish is a valuable model to understand the clinical onset and disease trajectories leading to serious clinical complications. While no morphological defects were observed during early larval stages in *tango2* mutants, a variability in survival rate was observed during larval and juvenile stages. During the first few days of development, zebrafish embryos and larvae survive on nutrients provided by the egg yolk. However, as these animals transition to external feeding at 5-6 days post fertilization, different amounts of nutrients and swimming (exercise) may elicit variable phenotypes in affected mutants. This variability in phenotype is also associated with *TANGO2* patients. Even the same genetic mutation has variable effects in different patients as a manifestation of genetic and variable environmental interactions (Lalani et al., 2016).

We further identified Tango2 is colocalized with SR, Golgi and mitochondrial markers in skeletal muscle. Previous studies have reported Tango2 localization in the endoplasmic reticulum (ER) and mitochondria in fibroblasts (Milev et al., 2021). These organelles are present in close proximity in the I-band region, therefore, there appears to be an overlap of Tango2 expression in these subcellular organelles. While high resolution imaging or in vivo proximity labeling based studies may provide a fine resolution spatial map of precise localization in skeletal muscle, ultrastructural defects in SR and mitochondria indicate the requirement of Tango2 function in these organelles. *tango2* mutants exhibit normal swimming behavior during the early larval stage (or the pre-symptomatic state) with10-15% of SR and mitochondria exhibiting ultrastructural defects in the muscle fibers. This suggests that tango2 deficiency results in intrinsic defects in SR and mitochondria which are still able to sustain basal threshold function in the skeletal muscle. However, under certain stress conditions, these defects prevent skeletal muscle to function beyond a basal threshold and finally result in muscle breakdown and other abnormalities.

ER is the primary hub of lipid biosynthesis and trafficking in cells. SR is the specialized form of ER that also regulates calcium handling in skeletal muscle to control excitation-contraction. SR stress induced by caffeine treatment resulted in an acute episode of motor deficits in *tango2* zebrafish mutants similar to episodic rhabdomyolysis seen in *TANGO2* patients. Caffeine binds to Ryr1 channels in SR and induces the release of Ca^2+^ in the cytosol. Dantrolene, a selective Ryr1 blocker was able to partially rescue caffeine induced motor function deficits in *tango2* mutants. However, a lack of complete rescue of motor function by dantrolene suggests that other processes could be contributing to motor function deficits in *tango2* mutants. Caffeine also increases intracellular levels of cyclic adenosine monophosphate (cAMP) by inhibiting phosphodiesterase enzymes and therefore resulting in lipolysis. Changes in the membrane phospholipids affect the activities of channel proteins that regulate calcium homeostasis in skeletal muscle as seen previously (Verkerke et al., 2019; Yamasaki et al., 2022). Thus, reduced abundance of phospholipids in Tango2 deficiency may impact channel activities and calcium homeostasis.

Previous studies have also reported the accumulation of fatty acids or acylcarnitines in TANGO2 patients derived cell lines. However, these findings are quite divergent and failed to provide a clear outcome due to wide clinical heterogeneity in patients samples. Our lipidomics analysis identified a significant reduction in phospholipids and triglycerides in *tango2* mutant zebrafish compared to controls. Phosphatidylcholine and phosphatidylethanolamine are the most abundant phospholipids (50% of total lipids) and a decrease in these lipid species may increase the susceptibility of membrane damage. Previous studies have shown a reduction in phospholipids resulted in skeletal muscle myopathy (Ferrara et al., 2021) and therefore, decreased amounts of phospholipids in tango2 deficiency could underlie the muscle weakness seen in *TANGO2* patients. Phospholipids synthesis occurs predominantly by the glycerol-3-phosphate pathway (Figure 5C). LPIN1 catalyzes an essential step of this process and mutations in *LPIN1* are the most common cause of severe recurrent rhabdomyolysis (Zeharia et al., 2008). These suggest that the glycerol-3-phosphate pathway is critical to disease pathogenesis in rhabdomyolysis. Glycerol-3 phosphate is also a substrate for triglyceride synthesis for energy storage. Reduced levels of triglycerides in *tango2* fish further point to defects in glycerolipid homeostasis in Tango2 deficiency. Some *TANGO2* patients also exhibit accumulation of acylcarnitine which suggests that defects in the glycerol-3-phosphate pathway in Tango2 deficiency may prevent utilization of acyl-CoA thus leading to acylcarnitine accumulation (Figure 5C). Acyl carnitine is normally metabolized by the ß-oxidation pathway. As mitochondrial defects are also observed in Tango2 deficiency (Figure 3), decreased metabolism of acylcarnitines may lead to its accumulation which is toxic for several organs, including the skeletal muscle, heart and liver. We did not observe any changes in the acylcarnitines in tango2 deficiency. As lipidomics analysis in *tango2* mutants was performed in the basal state, the accumulation of acylcarnitines seen in some *TANGO2* patients could be triggered by metabolic or other stress states. Taken together, our work demonstrates significant metabolic impairment in Tango2 deficiency in the absence of extrinsic triggers.

An accompanying study by Lujan et al. showed defects in lipid homeostasis on nutrient stress as a contributing factor in Tango2 disease pathogenesis through the regulation of acyl-CoA by phosphatidic acid (Lujan et al., 2022). In another accompanying study by Asadi et al., treatment with VitaminB5, a Coenzyme A precursor rescues seizures in a *Drosophila* model of Tango2 deficiency (Asadi, Milev, Saint-Dic, Gamberi, & Sacher, 2022). Therefore, these three studies point to perturbations in lipids biosynthesis and metabolism in Tango2 deficiency in different cellular and animal models. Future studies on how Tango2 regulates these processes will further improve our understanding of TANGO2-related disorders.

### Differences in *tango2* zebrafish models

A recent study showed that *TANGO2* orthologue *HRG-9* is critical for haem trafficking in *C. elegans* and deficiency of HRG-9 contributes to haem overload in mitochondria (Sun et al., 2022). Haem metabolism is critical for skeletal muscle function and mitochondria defects observed in our t*ango2* mutants may affect haem metabolism (Alves de Souza et al., 2021). However, a lack of difference in haem synthesis in control and *tango2* mutant fish in the previous study suggests TANGO2 is not required for *in vivo* haem metabolism in vertebrates (Sun et al., 2022). In addition, we found many differences between *tango2* mutants reported in the previous study and *tango2* mutants presented in this work. Sun et al., 2022 showed that *tango2* zebrafish mutants obtained from heterozygous mating were able to grow to adulthood with no obvious phenotype and display normal survival which was attributed to the extended stability of maternal mRNA. This contrasts with our findings which demonstrate *tango2* zebrafish mutants obtained from heterozygous matings exhibit reduced survival from the larval stage (80% at 6 dpf) to 4% by 3 months of age (n=350) (Figure 1D). Western blot analysis also failed to detect any Tango2 protein in our *tango2* mutants obtained from heterozygous crosses (Figure 1C) thus indicating the absence of maternal mRNA which is different than the previous study. Finally, the previous study showed that mutants obtained from mutant parents exhibited extensive defects in the brain, skeletal muscle and heart and die during the first two weeks of development. We also analyzed the *tango2* mutants obtained from *tango2* mutant parents in our zebrafish lines and did not see any defects in the brain, skeletal muscle and heart in the basal state as reported previously (Sun et al., 2022). Since different sgRNAs were used to create fish lines in ours and the previous study, we cannot rule out the possibility of some genetic background effect. Nonetheless, our results show that *tango2* mutants from either heterozygous or homozygous parents are morphologically and functionally similar, thus ruling out the maternal mRNA effect.

## Methods

### Zebrafish lines

Fish were bred and maintained using standard methods as described (Westerfield, 2000). All procedures were approved by the Brigham and Women’s Hospital Animal Care and Use Committee. *tango2*^*bwh210*^ *and tango2*^*bwh211*^ zebrafish lines were created in our laboratory by CRISPR-Cas9 approach. Zebrafish embryonic (0-2 days post fertilization) and larval stages (3-45 dpf), juvenile stage (45 dpf-3months) and adults (3 months) have been defined as described previously (Kimmel, Ballard, Kimmel, Ullmann, & Schilling, 1995). All studies presented in this work were performed on *tango2*^*bwh211*^ mutants obtained from heterozygous parents unless specified.

### Genotyping Assays for *tango2* lines

DNA was extracted from zebrafish larvae or fin-clips of adult zebrafish, genotyped by PCR and analyzed by 2% agarose gel (Bennett et al., 2018). PCR Primer sequences used for genotyping are 5’TGGGAATTAGCAAACGAGGA3’ and 5’ATGGCTGAAAGAGCTGTGCT3’.

### Western Blotting

Zebrafish larvae at 30 dpf were homogenized in buffer containing Tris–Cl (20 mM, pH 7.6), NaCl (50 mM), EDTA (1 mM), NP-40 (0.1%) and complete protease inhibitor cocktail (Roche Applied Sciences, Indianapolis, IN, USA). Following centrifugation at 11 000g at 4ºC for 15 min, protein concentration in supernatants was determined by BCA protein assay (Pierce, Rockford, IL, USA). Proteins were separated by electrophoresis on 4–12% gradient Tris–glycine gels (Invitrogen) and transferred onto polyvinylidene difluoride membrane using iblot dry blotting system (ThermoFisher Scientific).

Membranes were blocked in PBS containing 5% casein/0.1% Tween 20 and incubated with either rabbit polyclonal anti Tango2 (1:250, 27846-1-AP, Proteintech) or mouse monoclonal anti-*α*-tubulin (1:500, T9026-100UL, Sigma-Aldrich) primary antibodies. After washing, membranes were incubated with horseradish peroxidase-conjugated antirabbit (1: 2500, 170-6515) or antimouse (1: 5000, 170-6516) IgG secondary antibodies (BioRad, Hercules, CA, USA). Proteins were detected using the SuperSignal chemiluminescent substrate kit (Pierce).

### Myofiber isolation and immunofluorescence

Myofibers were isolated from control or *tnago2* larval zebrafish (45 dpf) as described previously with minor modifications (Ganassi, Zammit, & Hughes, 2021). Skinned zebrafish muscle samples were treated with collagenase for 90 minutes and triturated to release the myofibers. Myofibers were centrifuged at 1000g for 60 sec, washed and resuspended in DMEM media. Myofibers were plated on laminin coated 8 chamber permanox slides (Thermofisher Scientific) for further analysis. Fixed cells were blocked in 10% goat serum/0.3% Triton, incubated in primary antibody overnight at 4ºC, washed in PBS, incubated in secondary antibody for 1 h at room temperature (RT), washed in PBS, then mounted with Vectashield Mounting Medium (Vector Laboratories, Burlingame, CA, USA). Primary antibodies used were anti Tango2 (1:250, 27846-1-AP, Proteintech), mouse monoclonal anti sarcomeric *α*-actinin (1:100, A7732, Millipore Sigma), mouse monoclonal anti Ryr1 (1:100, R129-100UL, Millipore Sigma), anti Tomm20 (1:100, MABT166, Millipore Sigma) and Alexa fluor 568-phalloidin (1:100, Thermo Fisher Scientific, A12380). After washing in PBS several times, samples were incubated with anti-mouse Alexa Fluor (1:100, A-11005) secondary antibody (Thermofisher Scientific). Imaging was performed using a Nikon Ti2 spinning disk confocal microscope.

### Caffeine and Dantrolene treatment

Zebrafish (6dpf) obtained from heterozygous mattings were placed in individual wells of a 48 well dish and swimming behavior was analyzed at the basal level. Caffeine and dantrolene treatment were performed as previously described with some modifications (Endo et al., 2022). Subsequently, E3 water was replaced with 0.5uM caffeine containing E3 water and fish were incubated for 1 hr and caffeine was replaced with normal E3 water. Swimming behavior was analyzed again after an hour of recovery and after 24 hrs of recovery by the automated tracking system. For dantrolene (Millipore Sigma D9175) treatment, zebrafish were incubated with 5uM dantrolene for 2 hrs followed by 1 hr incubation with caffeine and dantrolene. Swimming behavior was analyzed before the treatment, after an hour of recovery and after 24 hrs of recovery by automated tracking system.

### Lipidomic Profiling

Control and *tango2* mutant zebrafish (4 weeks, n=5 each) were homogenized with 1 ml of MBTE. 300uL of methanol with internal standard was added and samples were mixed for 10 mins. 200 uL of water was added to facilitate phase separation. The extracts were centrifuged at 20,000 rcf for 10 mins. The top layer was removed, dried and reconstituted in 150ul of IPA for analysis. Analysis was performed using a Thermo Q Exactive Plus coupled to a Waters Acquity H-Class LC. A 100 mm x 2.1 mm, 2.1 µm Waters BEH C18 column was used for separations. The following mobile phases were used: A-60/40 ACN/H20 B-90/10 IPA/ACN; both mobile phases contained 10 mM Ammonium Formate and 0.1% Formic Acid. A flow rate of 0.2 mL/min was used. Starting composition was 32% B, which increased to 40% B at 1 min (held until 1.5 min) then 45% B at 4 minutes. This was increased to 50% B at 5 min, 60% B at 8 min, 70% B at 11 min, and 80% B at 14 min (held until 16 min). At 16 min the composition switched back to starting conditions (32% B) and was held for 4 min to re-equilibrate the column. Samples were analyzed in positive/negative switching ionization mode with top 5 data dependent fragmentation.

Raw data were analyzed by LipidSearch. Lipids were identified by MS2 fragmentation (mass error of precursor=5 ppm, mass error of product=8 ppm). The identifications were generated individually for each sample and then aligned by grouping the samples (OxPAPC=C, HF=S1, Con=S2). Normalization was performed using EquiSplash from Avanti. Samples were normalized and biological replicates were averaged. P-value and fold change were calculated as instructed as previously described (Aguilan, Kulej, & Sidoli, 2020). P-value was set to 0.05.

### Zebrafish locomotion assay

Zebrafish swimming behavior was quantified by using the Zantiks MWP automated tracking system (Zantiks Ltd., Cambridge, UK). Larval zebrafish (5-7 dpf) were placed individually into each well of a 48 well plate and their swimming behavior was recorded for 50 minutes (10 minutes light, 10 minutes dark, 10 minutes light, 10 minutes dark, 10 minutes light, end). Four independent blind trials were performed, and total distance and cumulative duration of the movement were recorded.

### Electron Microscopy

7 dpf zebrafish embryos were used to perform transmission electron microscopy. Heads of individual larval fish were removed for genotyping and bodies were fixed in formaldehyde–glutaraldehyde–picric acid in cacodylate buffer overnight at 4ºC, followed by osmication and uranyl acetate staining. Subsequently, embryos were dehydrated in a series of ethanol washes and embedded in TAAB Epon (Marivac Ltd., Halifax, NS, Canada). Sections (95 nm) were cut with a Leica UltraCut microtome, picked up on 100 mm Formvar-coated copper grids, and stained with 0.2% lead citrate. Sections were viewed and imaged using a JEOL 1200EX transmission electron microscope at the Harvard Medical School Electron Microscopy Core.

### Quantification and statistical analysis

All samples were blinded till final analyses and statistical analyses were performed using GraphPad Prism9.

## Acknowledgements

VAG was supported by NIH R56AR077017 and A Foundation Building Strength grant. We thank Dr. Behzad Moghadaszadeh for the helpful discussions. Authors also thank Louis Trakimas and Anja Nordstrom at the electron microscopy core (Harvard Medical School) for assistance with sample preparation.

